# The unconventional kinesin Kif26a is required for the guidance of major axon tracts in the developing mouse brain

**DOI:** 10.64898/2026.05.20.726728

**Authors:** Srisathya Srinivasan, Kayla Louie, Timothy Casey-Clyde, Zixin Wei, Morgan Krueger, Edith Karuna, Ryuichi Nishinakamura, Konstantinos Zarbalis, Qizhi Gong, Sergi Simo, Hsin-Yi Henry Ho

**Affiliations:** Department of Cell Biology and Human Anatomy, School of Medicine, University of California, Davis, California, United States; Department of Pathology and Laboratory Medicine, School of Medicine, University of California, Davis, United States; Department of Kidney Development, Institute of Molecular Embryology and Genetics, Kumamoto University, Kumamoto, Japan

**Author notes:** Co-first authors (Srisathya Srinivasan, Kayla Louie, Timothy Casey-Clyde).

## Abstract

*Kif26a* and *Kif26b* encode a family of unconventional kinesins with emerging roles in neurodevelopment, and mutations in both genes have been implicated in a spectrum of neurodevelopmental disorders. The precise mechanisms by which the Kif26 family orchestrates mammalian brain development, however, remain unclear. In this study, we show that *Kif26a* and *Kif26b* are expressed in distinct regions of the developing mouse brain. Using a new allelic series in mice, we demonstrate that *Kif26a* is required for the guidance of multiple forebrain axon tracts. This requirement is direct and cell autonomous, as cell proliferation, survival, and cortical layering are unaffected in the *Kif26a* mutants. These guidance defects closely resemble those reported for the Fzd3-Celsr3-Dystroglycan pathway, suggesting that Kif26a may participate within this conserved signaling axis to steer growing axons.

## Introduction

Nervous system development requires the precise coordination of many cellular events. A central goal of modern neurobiology is to identify the molecules and pathways that orchestrate these processes and understand how their dysfunction causes disease. The Kif26 family of kinesin motor proteins has long been linked to nervous system development. *Vab-8*, the sole homologue in *C. elegans*, was first identified more than 30 years ago as a key regulator of neuronal migration and axon guidance (Wightman et al., 1996, Wolf et al., 1998). *Vab-8* mutants display specific defects in the posteriorly directed movements of multiple neuronal and axonal populations, and a subset of these phenotypes is shared by mutants in the Wnt/Frizzled pathway (Zinovyeva et al., 2008, Pan et al., 2006, Jain and Lundquist, 2025). The vertebrate genome encodes two related family members, *Kif26a* and *Kif26b* (Miki et al., 2001). Unlike conventional kinesins, both proteins are thought to lack the ATPase activity typically associated with motor function and are therefore classified as unconventional kinesins (Zhou et al., 2009, Hirokawa et al., 2009). Although the molecular functions of the family remain poorly defined, studies in mice and cultured cells have shown that Kif26a is required for enteric nervous system development, while Kif26b acts prominently in the developing kidney, gonad, vascular system (Zhou et al., 2009, Uchiyama et al., 2010, Guillabert-Gourgues et al., 2016, Susman et al., 2017). Moreover, Kif26b functions in noncanonical Wnt pathways to regulate cell migration, polarity, and adhesion (Susman et al., 2017, Guillabert-Gourgues et al., 2016).

Human genetics has revealed a crucial role for the Kif26 family in neurodevelopmental disorders. The first disease-associated variant identified was a missense mutation in *KIF26B* (G546S), mapped in an infant with pontocerebellar hypoplasia (Wojcik et al., 2018); a second *KIF26B* variant (D1904N) was later reported in patients with spinocerebellar ataxia (Nibbeling et al., 2017). More recently, homozygous loss-of-function mutations in *KIF26A* have been reported in patient cohorts with severe brain malformations, as well as in separate cohorts presenting with hydrocephalus, megacolon and/or cranial dysinnervation disorder (Qian et al., 2022, Almannai et al., 2023, Akula et al., 2023, Gregg et al., 2024). Together with the earlier work on *Vab-8* in *C. elegans*, these findings underscore the importance of the Kif26 family in mammalian nervous system development and highlight the conservation of its function across metazoans.

The mechanisms by which the Kif26 family mediates brain development, however, remain incompletely understood. No studies to date have used an intact animal model to investigate the patterns of *Kif26a* and *Kif26b* expression during mammalian brain development. Likewise, the *in vivo* requirements of these genes, as well as their potential functional redundancy, have not been systematically examined *in vivo*. Although RNAi experiments in cultured neurons, organoids, and *in utero* electroporation models have shown that loss of *Kif26a* expression can perturb proliferation, survival, and radial migration during corticogenesis (Qian et al., 2022), the effects of germline *Kif26a* ablation on these processes, and whether *Kif26a* has additional functions during brain development, remain to be determined.

Here, we report the generation and characterization of a new *Kif26a* allelic series in mice - a *LacZ* expression reporter, a constitutive null, and a conditional allele - that together allow *Kif26a* expression and loss-of-function phenotypes to be studied *in vivo*. Using the reporter line, along with RNAscope and single-cell transcriptomic analyses, we map the distinct expression domains of *Kif26a* and *Kif26b* in the developing brain. Phenotypic analyses of the mutants reveal a specific, cell autonomous requirement for *Kif26a* in the guidance of major forebrain axon tracts. These guidance phenotypes closely parallel those reported for the Fzd3-Celsr3-Dystroglycan pathway, suggesting that *Kif26a* may act within this signaling axis to direct axon pathfinding.

## Results

### *Kif26a* and *Kif26b* are expressed in distinct regions of the developing mouse brain

To gain insight into the functions of *Kif26a* and *Kif26b*, we first examined their expression patterns by RNAscope. At midgestation (E14.5), *Kif26a* is expressed prominently in the cortical plate and striatum, and at a lower level in the piriform cortex, the septal neuroepithelium, and the ventricular/subventricular zones (VZ/SVZ) (**Fig. 1A**). At E18.5, *Kif26a* remains strongly expressed in the deeper layers (layers V and VI) and moderately in the striatum, while its signal in the VZ/SVZ has largely diminished **(Fig. 1C**). Some expression is also observed around the nucleus accumbens (**Fig. 1C**).

**Figure 1.**
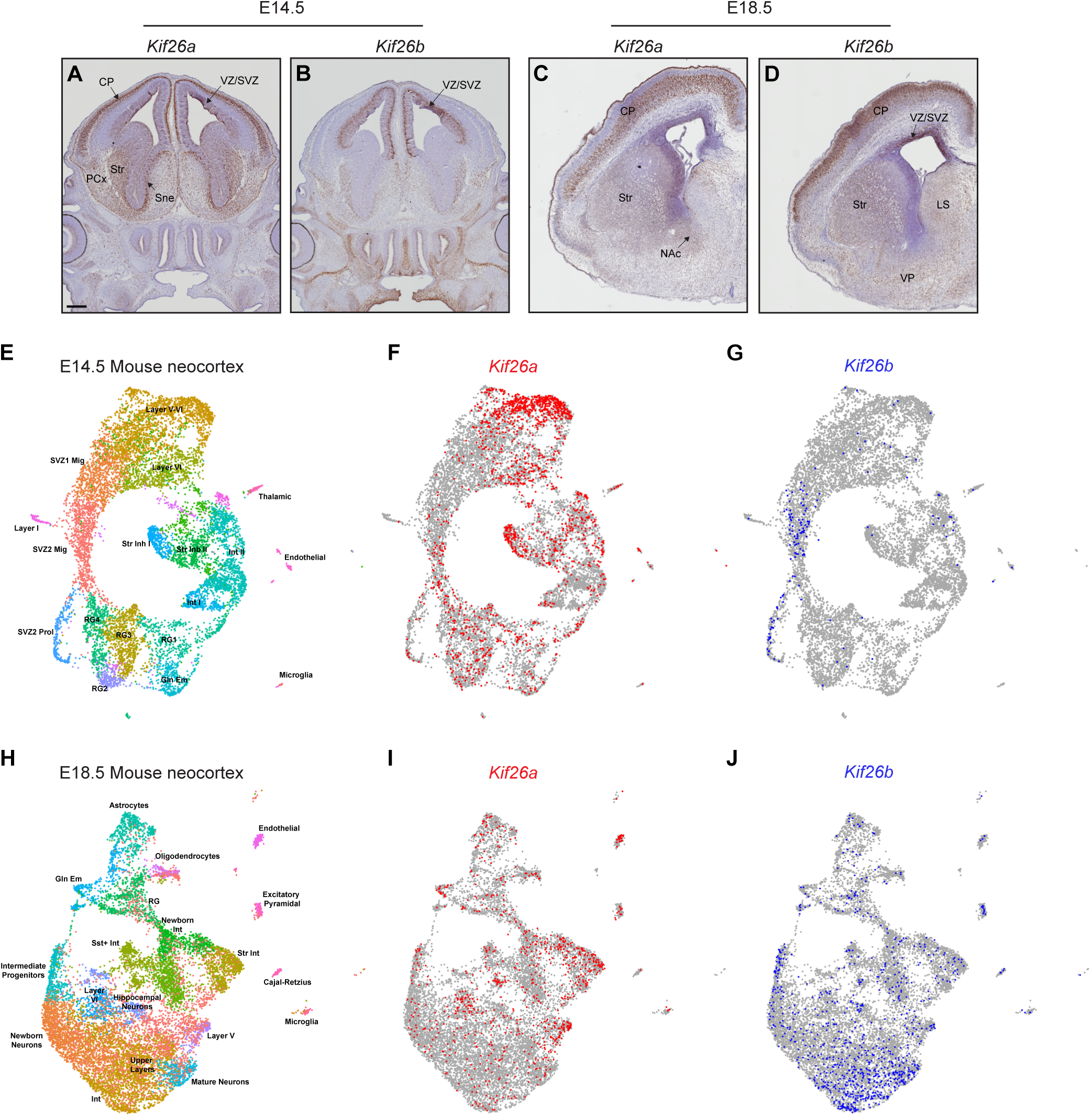
Spatiotemporal expression of *Kif26a* and *Kif26b* in the developing mouse brain. (**A-D**) Expression of *Kif26a* and *Kif26b* mRNA in different brain regions at E14.5 (**A, B**) and E18.5 (**C, D**), as determined by RNAscope-based in situ hybridization. Scale bar: 250 µm. (**E-J**) Expression of *Kif26a* and *Kif26b* mRNA in distinct cell populations of the developing neocortex at E14.5 (**E-G**) and E18.5 (**H-J**), as determined by analysis of publicly available single-cell RNA sequencing data (Loo et al., 2019, Wittmann et al., 2021). Processed data are shown as UMAP plots of cell clusters with *Kif26a*- (**F, I**) and *Kif26b-* (**G, J**) expressing cells shown in red and blue, respectively. CP, cortical plate; Str, striatum or striatal; VZ, ventricular zone; SVZ, sub ventricular zone; PCx, piriform cortex; Sne, septal neuroepithelium; NAc, nucleus accumbens; LS, lateral septal nucleus; VP, ventral pallidum; Mig, migrating; Prol, proliferating; Int, interneurons; Inh, inhibitory; RG, radial glia; Gln Em, ganglionic eminence; Sst+, somatostatin positive.

In contrast, *Kif26b* shows a largely distinct expression pattern. At E14.5, *Kif26b* is expressed prominently in the VZ/SVZ, particularly the SVZ, moderately in the lateral aspect of the striatum, but absent in the cortical plate (**Fig. 1B**). By E18.5, *Kif26b* expression has broadened to include the upper cortical layers and the striatum in addition to the VZ/SVZ, with diffuse *Kif26b* signal in the lateral septal nucleus and ventral pallidum (**Fig. 1D**).

Analysis of publicly available single-cell transcriptomic data (Loo et al., 2019, Wittmann et al., 2021) corroborates the RNAscope findings (**Fig. 1E-J**). At E14.5, *Kif26a* is again most strongly expressed in layer V and VI cortical neurons and in the striatum, whereas *Kif26b* is largely restricted to progenitor populations in the SVZ (**Fig. 1E, 1F**). At E18.5, *Kif26a* remains enriched in deep cortical layers and striatal interneurons but is now also present in a subset of pyramidal neurons and endothelial cells (**Fig. 1H, 1I**). In contrast, *Kif26b* expression broadens to include upper-layer cortical and other mature neuronal populations, as well as intermediate progenitors (**Fig. 1H, 1J**). Together, these data provide the first comparative analysis of *Kif26a* and *Kif26b* expression in the developing mammalian brain, and the distinct spatiotemporal profiles suggest that the two genes play non-redundant roles during nervous system development.

### Generation of a *Kif26a* mutant allelic series in mice for *in vivo* expression and functional studies

To define the *in vivo* requirements of the Kif26 family during brain development, we sought to examine mutant mice for phenotypes. A constitutive *Kif26a* knockout strain, in which exon 7 was deleted, has been described previously (Zhou et al., 2009). The mutant pups die within ∼15 days of birth from megacolon and growth retardation, but no brain phenotypes were reported (Zhou et al., 2009). We were unable to obtain the line for further analysis.

We therefore generated a new *Kif26a* allelic series in mice using the “knockout-first” strategy described by Skarnes *et al*. (Skarnes et al., 2011). Homologous recombination in embryonic stem (ES) cells was used to introduce a gene-trap cassette containing: 1) a splice acceptor, 2) an *IRES-LacZ* transcriptional reporter, 3) a transcriptional termination (*STOP*) signal containing the SV40 polyadenylation sequence, and 4) a neomycin resistance gene into the intron preceding exon 4 of the *Kif26a* gene (**Fig. 2A**). Two *frt* and three *lox2272* sites were also positioned such that conditional and constitutive knockout alleles could be derived from the founder (**Fig. 2A**). Exon 4 was selected because its excision is predicted to introduce an early frameshift that disrupts 87.4% of the *Kif26a* open reading frame. Targeted ES cells were injected into donor blastocysts to generate chimeras. Crossing of the founders to a germline Cre driver (*Sox2^Cre^*) excised both the neomycin-resistance cassette and exon 4, giving rise to the *Kif26a^LacZ^* allele (**Fig. 2A**). Because *LacZ* in this allele is driven by the endogenous *Kif26a* promoter, *Kif26a^LacZ^* functions as a transcriptional reporter. X-gal staining of E14.5 *Kif26a^+/LacZ^*brains revealed expression patterns closely matching the RNAscope data (**Fig 2B, 2D, 2D’**). No staining was observed in wild-type (WT) control brains (**Fig 2C, 2E, 2E’**), confirming the specificity of the signal.

**Figure 2.**
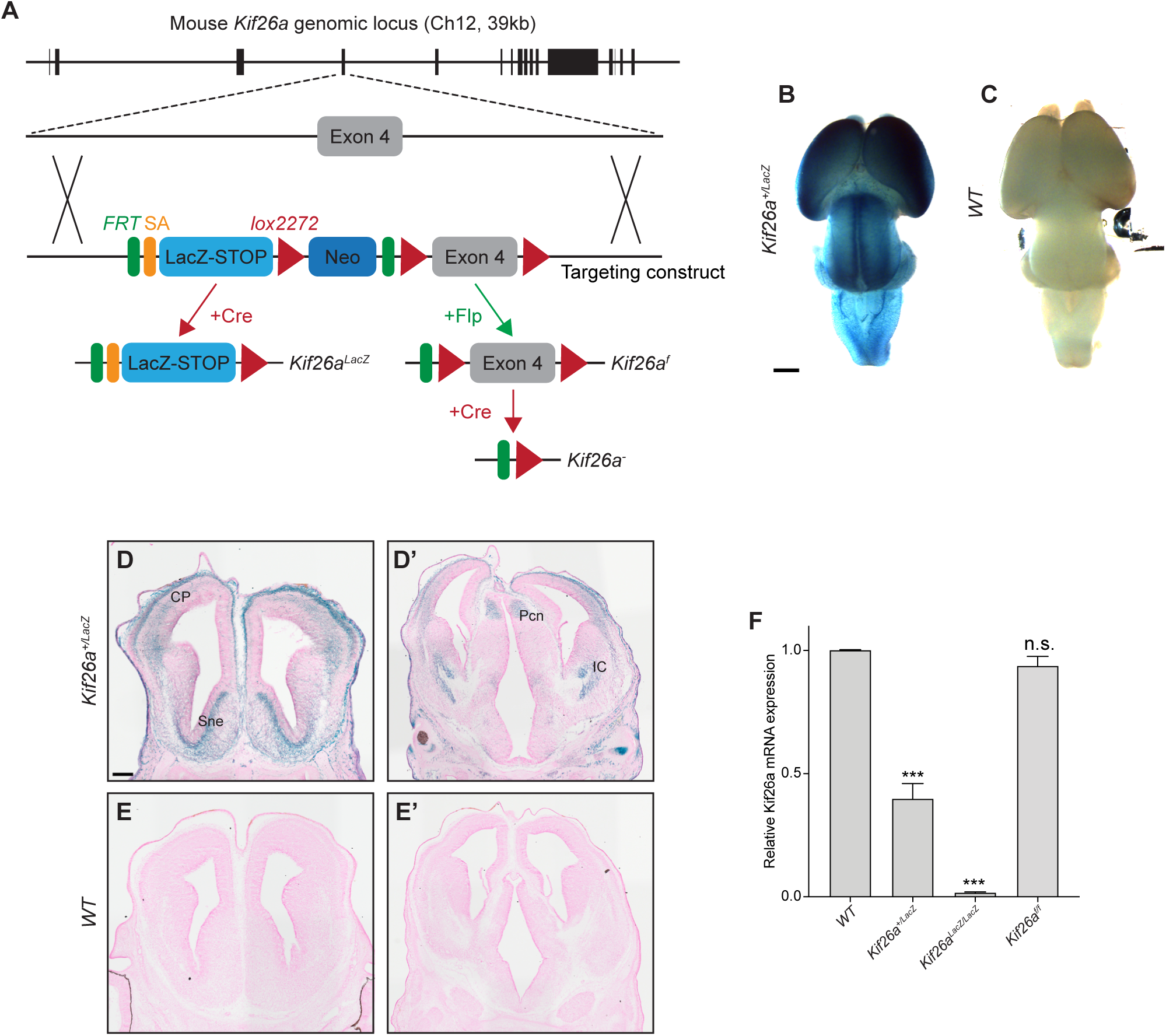
Genetic modifications of the mouse Kif26a locus. (**A**) Schematic of the *Kif26a* gene targeting strategy. A multifunctional allele of *Kif26a* was generated by homologous recombination in mouse embryonic stem (ES) cells. A splice acceptor (SA)-LacZ-STOP cassette, a neomycin (Neo) resistance gene, two *frt* and three *lox2272* sites were introduced into the intronic regions surrounding exon 4 of the *Kif26a* locus on chromosome 12 (Ch 12). Modified ES cells were injected into donor blastocysts to derive the founder strain. Crossing the founder mice with a germline-specific Cre (*Sox2^Cre^*) driver generates the *Kif26a^LacZ^* allele. The insertion of SA-LacZ allows *Kif26a^LacZ^* to function as a transcription reporter, while the STOP element renders this allele transcription null. Crossing of the founder mice with a germline-specific Flp (*ROSA26-FLPe*) driver gave rise to the conditional *Kif26a^f^* allele which can further undergo Cre-mediated recombination (e.g. using tissue-specific Cre drivers) to generate the *Kif26a^−^* allele. (**B-E’**) Validating the reporter function of *Kif26a^LacZ^*. Wholemount X-gal staining of the E14.5 *Kif26a^+/LacZ^* brain shows widespread X-gal signals (blue) **(B)**. X-gal reactivity is not present in the wild-type (WT) control brain (**C**), confirming that the signals observed in the *Kif26a^+/LacZ^* brains are specific. Scale bar: 1 mm. X-gal staining of E14.5 coronal series from rostral (**D**) and caudal (**D’**) regions of the *Kif26a^+/LacZ^* brain shows prominent X-gal signals in distinct cortical and subcortical structures, including the cortical plate (CP), septal neuroepithelium (Sne), precommissural nucleus (PCN) and internal capsule (IC). X-gal reactivity is not present in the WT control sections (**E** and **E’**, rostral and caudal sections, respectively). Scale bar: 250 µm. **(F)** Reverse transcription and quantitative PCR (qRT-PCR) verified the absence of *Kif26a* mRNA in *Kif26a^LacZ/LacZ^*animals. *Kif26a* expression in *Kif26a^f/f^* animals is indistinguishable from that of the WT control. Relative *Kif26a* mRNA expression in the mutant groups is compared to that of WT. Error bars represent +SEM calculated from three independent biological samples (embryos for each genotype). qPCR reactions were performed in three technical replicates for each biological sample. t-Test (unpaired) was performed to determine statistical significance of each mutant sample vs. *WT*. ***, *p* < 0.001; n.s., not significant.

The transcriptional termination signal in the *LacZ-STOP* cassette, together with the excision of exon 4, predicts *Kif26a^LacZ^* to be a functionally null allele. RT-qPCR confirmed that no *Kif26a* transcript was detectable in homozygous *Kif26a^LacZ/LacZ^* embryos (**Fig 2F**). Heterozygous *Kif26a^+/LacZ^* animals are viable, fertile and indistinguishable from their WT littermates, whereas homozygous *Kif26a^LacZ/LacZ^* mice die within a few hours of birth. Loss of *Kif26a* expression therefore causes recessive perinatal lethality.

To generate the conditional *Kif26a^f^* allele, we crossed the founder line to a germline Flp driver (*Rosa26-FlpE*). *Kif26a^f/f^* mice exhibit normal levels of *Kif26a* expression and are phenotypically indistinguishable from WT (**Fig 2F**). Subsequent crossing to *Sox2^Cre^* produced the *Kif26a^−^* allele (**Fig 2A)**. Next-generation sequencing of cDNA from *Kif26a^−/-^*embryos confirmed the expected frameshift from exon 4 excision. Approximately 80% of *Kif26a^−/-^* animals die perinatally, with the majority of surviving pups developing megacolon, consistent with the previous report (Zhou et al., 2009).

Given that the *Kif26a^LacZ^*allele eliminates *Kif26a* transcription and exhibits a more penetrant perinatal lethality phenotype than *Kif26a^−^*, it was used as the primary germline knockout model for subsequent phenotypic analyses.

### *Kif26a* is required for the formation of major axon tracts in the developing forebrain

Given the severe neurodevelopmental phenotypes in patients with *KIF26A* mutations and the strong expression of *Kif26a* in the developing mouse cortex, we analyzed *Kif26a^LacZ/LacZ^* brains for possible phenotypes. At E18.5, all major brain regions – cerebral hemispheres, olfactory bulbs, thalami, midbrain, pons, medulla, and cerebellum – are formed. However, the cerebral cortices of *Kif26a^LacZ/LacZ^* mice appear partially collapsed, likely due to thinning of the underlying structures (**Fig. 3B, B’**). Indeed, L1-CAM staining of coronal sections revealed pronounced defects in several major fiber tracts, including the complete loss of the anterior commissure, a substantial reduction in the number and cross-sectional area of internal capsule fibers, and thinning of the intermediate zone (**Fig. 3B, 3B’, 3E-I**). These fiber tract phenotypes have not been previously described in either patients or animal models. The defects are absent in *Kif26b* single mutants (**Fig. 3C, 3C’, 3E-I**) but present in *Kif26a^LacZ/LacZ^; Kif26b^−/-^* double mutants (**Fig. 3D, 3D’; 3E-I**), indicating that *Kif26b* does not compensate for the loss of *Kif26a* in this context. We therefore focused the remainder of the study on determining the developmental mechanisms underlying these defects. Curiously, corpus callosum agenesis – a hallmark of patients carrying *KIF26A* loss-of-function mutations (Qian et al., 2022) – is not observed in either *Kif26a^LacZ/LacZ^* single or *Kif26a^LacZ/LacZ^; Kif26b^−/-^* double mutant brains (**Fig. 3A-D’, 3I**). We will return to this difference between mouse and human phenotypes in the Discussion.

**Figure 3.**
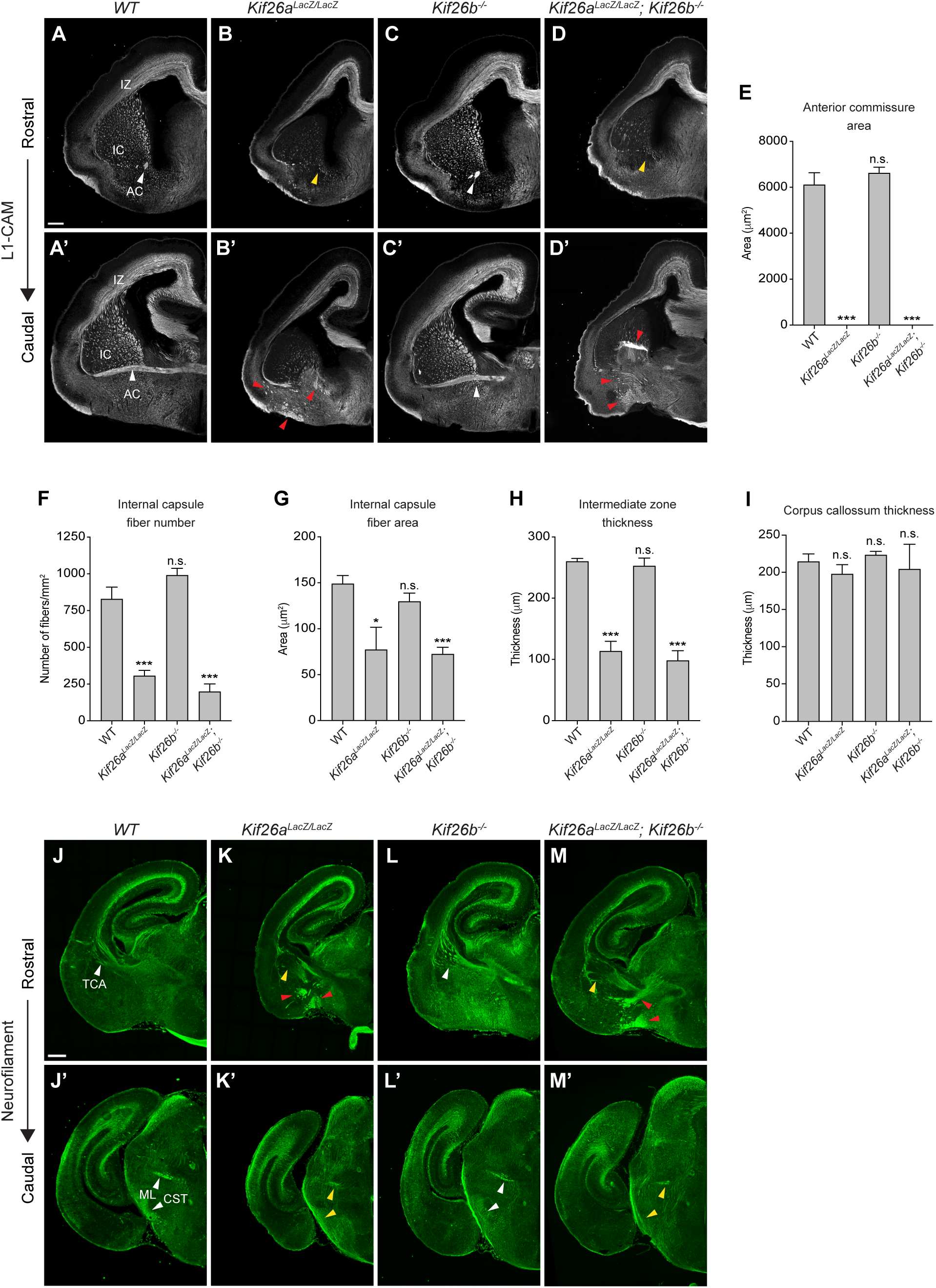
Loss of major fiber tracts in the *Kif26a^LacZ/LacZ^ brain*. (**A-D’**) L1-CAM immunostaining of E18.5 coronal series showing normal (white arrow heads) and absent (yellow arrow heads) fiber tracts in WT (**A, A’**), *Kif26a^LacZ/LacZ^* (**B, B’**), *Kif26b^−/-^* (**C, C’**), and *Kif26a^LacZ/LacZ^; Kif26b^−/-^* (**D, D’**) brains. Affected axon tracts include the anterior commissure (AC), fibers passing through the internal capsule (IC) and fibers in the intermediate zone (IZ). Aberrant axons are visible in *Kif26a^LacZ/LacZ^*and *Kif26a^LacZ/LacZ^; Kif26b^−/-^* brains (denoted by the red arrow heads). Scale bar: 250 µm. (**E-I**), quantification of the fiber tract phenotypes shown in **A-D’**. Data are represented as mean +SEM of the quantified phenotypes from 3 independent embryos for each genotype. t-Test (unpaired) was performed to determine statistical significance of mutants vs. *WT*. ***, *p* < 0.001; *, *p* < 0.05; n.s., not significant. (**J-M’**) Neurofilament immunostaining of E18.5 coronal series showing normal (white arrow heads) and absent (yellow arrow heads) fiber tracts in WT (**J, J’**), *Kif26a^LacZ/LacZ^* (**K, K’**), *Kif26b^−/-^* (**L, L’**), and *Kif26a^LacZ/LacZ^; Kif26b^−/-^* (**M, M’**) brains. Affected axon tracts include the thalamocortical axons (TCA), medial lemniscus (ML), and corticospinal tract (CST). Aberrant fibers are indicated by the red arrowheads (**K, M**). Scale bar: 250 µm.

Closer examination of the L1-CAM-stained sections revealed ectopic fibers in tissues surrounding the striatum in *Kif26a^LacZ/LacZ^* and *Kif26a^LacZ/LacZ^; Kif26b^−/-^* brains (**Fig. 3B’, 3D**’). The location and trajectory of these fibers are suggestive of misrouted thalamocortical axons (TCA) and corticothalamic (CTA) axons and hint at a role for Kif26a in axon pathfinding. Together with the compromised anterior commissure, internal capsule, and intermediate zone, these phenotypes closely resemble the axon guidance defects reported in mice lacking Frizzled-3 (Fzd3), Celsr3 and Dystroglycan (Wang et al., 2002, Lyuksyutova et al., 2003, Tissir et al., 2005, Zhou et al., 2008, van Amerongen et al., 2012, Hua et al., 2014, Qu et al., 2014, Lindenmaier et al., 2019). Fzd3 is a Frizzled Wnt receptor involved in planar cell polarity (PCP) signaling; Celsr3 is a cadherin family receptor that also functions in PCP; and Dystroglycan is a dystrophin associated glycoprotein that binds Celsr3 to regulate neuronal adhesion (Wang et al., 2006, Butler and Wallingford, 2017, Wang et al., 2016, Chai et al., 2015, Lindenmaier et al., 2019). These three molecules act in a common pathway to control axon guidance, and we hypothesize that Kif26a may also be a part of this signaling axis.

To test whether the *Kif26a^LacZ/LacZ^* brain shares additional phenotypes with *Fzd3*, *Celsr3*, or *Dystroglycan* mutants, we performed neurofilament staining to survey a broader set of fiber tracts. This confirmed the aberrant fibers associated with the CTA and TCA observed by L1-CAM staining (**Fig. 3K, 3M**) and uncovered further defects in the corticospinal tract and medial lemniscus (**Fig. 3K’, 3M’**), similar to the defects reported in the *Fzd3* mutant (Wang et al., 2002).

RNAi-based perturbation of *Kif26a* in mouse *in utero* electroporation and human organoid models have been reported to cause corticogenesis defects and neuronal apoptosis (Qian et al., 2022), which could in principle affect axon formation secondarily. To test whether such defects are present in knockout mice, we stained *Kif26a^LacZ/LacZ^* cortices for layer specific markers (SATB2 for layers II-IV; CTIP2 for layers V and VI; TBR1 for layer VI and the subplate) and found no statistically significant difference in lamination from WT (**Fig. 3 supplement 1A, B**). The SATB2+, CTIP2+ and, TBR1+ layers were slightly thinner in the mutants (**Fig. 3 supplement 1C**), but neuronal density within each layer was unchanged (**Fig. 3 supplement 1D**). EdU incorporation and TUNEL assays similarly revealed no significant differences in cell proliferation (**Fig. 3 supplement 1E, 1F**) or apoptosis (**Fig. 3 supplement 1G, 1H**). We conclude that the fiber tract defects are unlikely to arise secondarily from earlier deficits in proliferation, survival, or cortical lamination, and that Kif26a’s role in axon development is direct.

**Figure 3 supplement 1.**
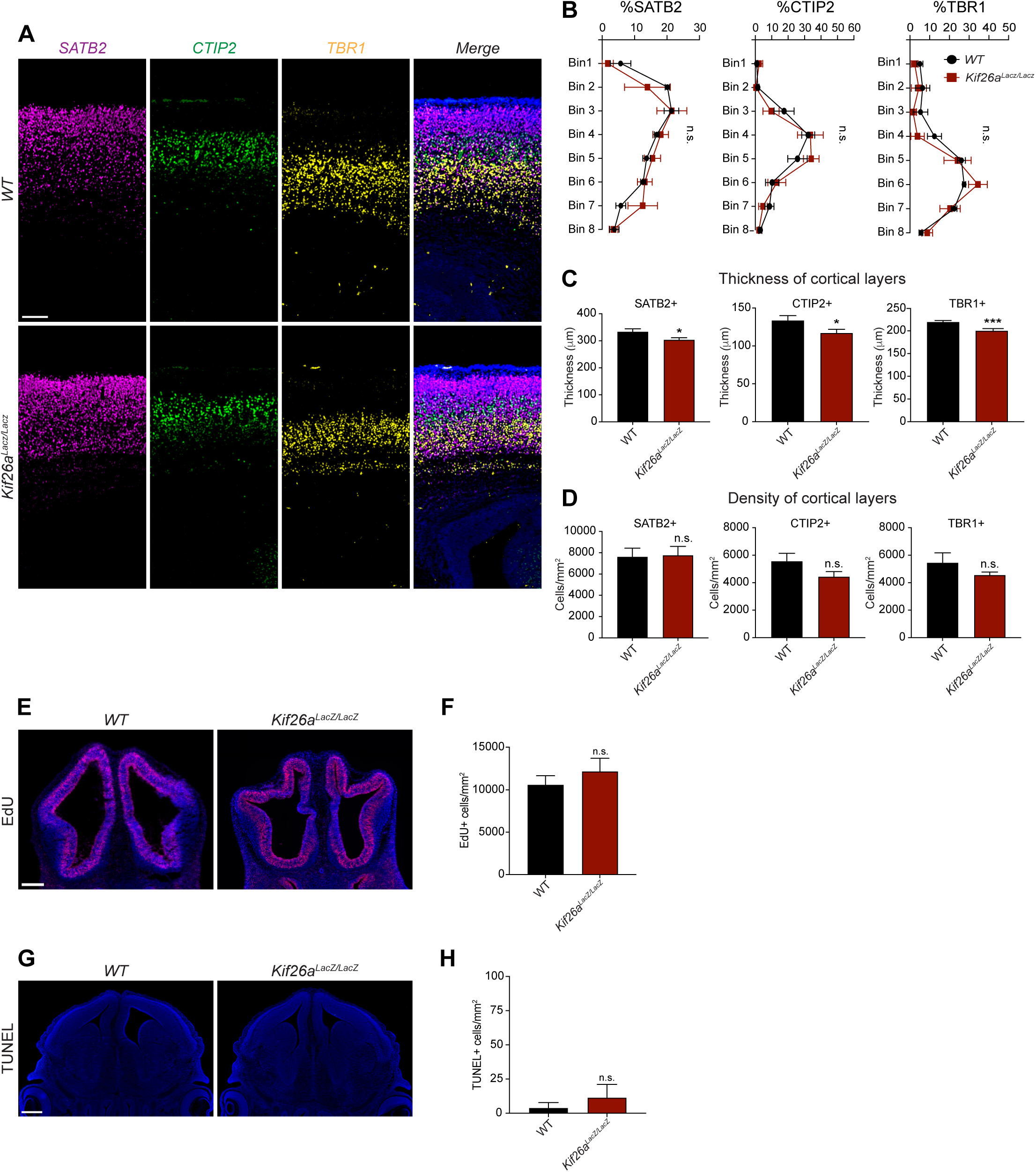
Analysis of cortical layering, cell proliferation and apoptosis in the *Kif26a^LacZ/LacZ^ brain*. **(A)** E18.5 cortical sections were immunostained with anti-SATB2 (magenta), -CTIP2 (green), and -TBR1 (yellow) antibodies to mark the superficial, intermediate, and deep cortical layers, respectively. Sections were counterstained with DAPI (blue). Scale bar: 100 µm. **(B)** Distribution of SATB2^+^, CTIP2^+^ and TBR1^+^ cells across the thickness of WT and *Kif26a^LacZ/LacZ^* cortices. The region encompassing the marginal zone to the subplate was virtually divided into 8 bins, and the percentage of cells positive for each marker was quantified for each bin. **(C)** Quantification of the thickness of SATB2^+^, CTIP2^+^, and TBR1^+^ layers in WT and *Kif26a^LacZ/LacZ^* brains. **(D)** Quantification of the density of SATB2^+^, CTIP2^+^, and TBR1^+^ neurons within the respective layers of the WT and *Kif26a^LacZ/LacZ^* brains. **(E)** Coronal sections of E12.5 WT and *Kif26a^LacZ/LacZ^* brains showing the incorporation of EdU (red) as a proxy of cell proliferation. EdU was injected intraperitoneally into pregnant dams carrying E12.5 embryos. Embryos were collected 4 hr post injection. Sections were counterstained with Hoechst 33342 (blue). Scale bar: 250 µm. (**F**) Quantification of EdU incorporation (number of EdU^+^ cells per mm^2^) in the cortex of WT and *Kif26a^LacZ/Lac^*^Z^ brains. (**G**) Coronal sections of E14.5 WT and *Kif26a^LacZ/LacZ^* brains analyzed for apoptosis by TUNEL assay. Apoptotic cells are shown in green. Sections were counterstained with Hoechst 33342 (blue). Scale bar: 250 µm. (**H**) Quantification of apoptosis (number of TUNEL^+^ cells per mm^2^) in WT and *Kif26a^LacZ/LacZ^* brains. Data are represented as mean +SEM of the quantified phenotypes from 3 independent embryos for each genotype. t-Test (unpaired) was performed to determine statistical significance of mutant vs. WT. ***, *p* < 0.001; *, *p* < 0.05; n.s., not significant.

### Anterograde axon tracing reveals a functional role of Kif26a in axon guidance

To directly test whether loss of Kif26a causes axon misguidance, we performed anterograde axon tracing using the lipophilic dye DiI (1,1’-dioctadecyl-3,3,3’,3’-tetramethylindocarbocyanine perchlorate). Given the appearance of aberrant fibers around the striatum in *Kif26a^LacZ/LacZ^* brains (**Fig. 3B’, 3D’, 3K, 3M**), we focused our initial analysis on the CTA and TCA. In WT brains, DiI crystals placed in the dorsal cortex labeled the expected CTA trajectory (**Fig. 4A, 4A’**): fibers exit the cortex via the intermediate zone, turn ventrally toward the pallial-subpallial boundary (PSPB), course medially through the striatum, and cross the diencephalon-telencephalon boundary (DTB) to enter the dorsal thalamus. In *Kif26a^LacZ/LacZ^* brains, descending CTA fibers stalled at the PSPB and frequently misrouted along the external capsule (**Fig. 4B, 4B’**). The reciprocal experiment showed similar defects: WT TCA axons crossed the DTB and the striatum and turned toward the PSPB to enter the cortex (**Fig. 4C, 4C’**), whereas mutant TCA axons stalled at the DTB and then extended ventrally out of the diencephalon without turning (**Fig. 4D, 4D’**). Together, these tracing experiments demonstrate a direct role for Kif26a in guiding the CTA and TCA axons.

**Figure 4.**
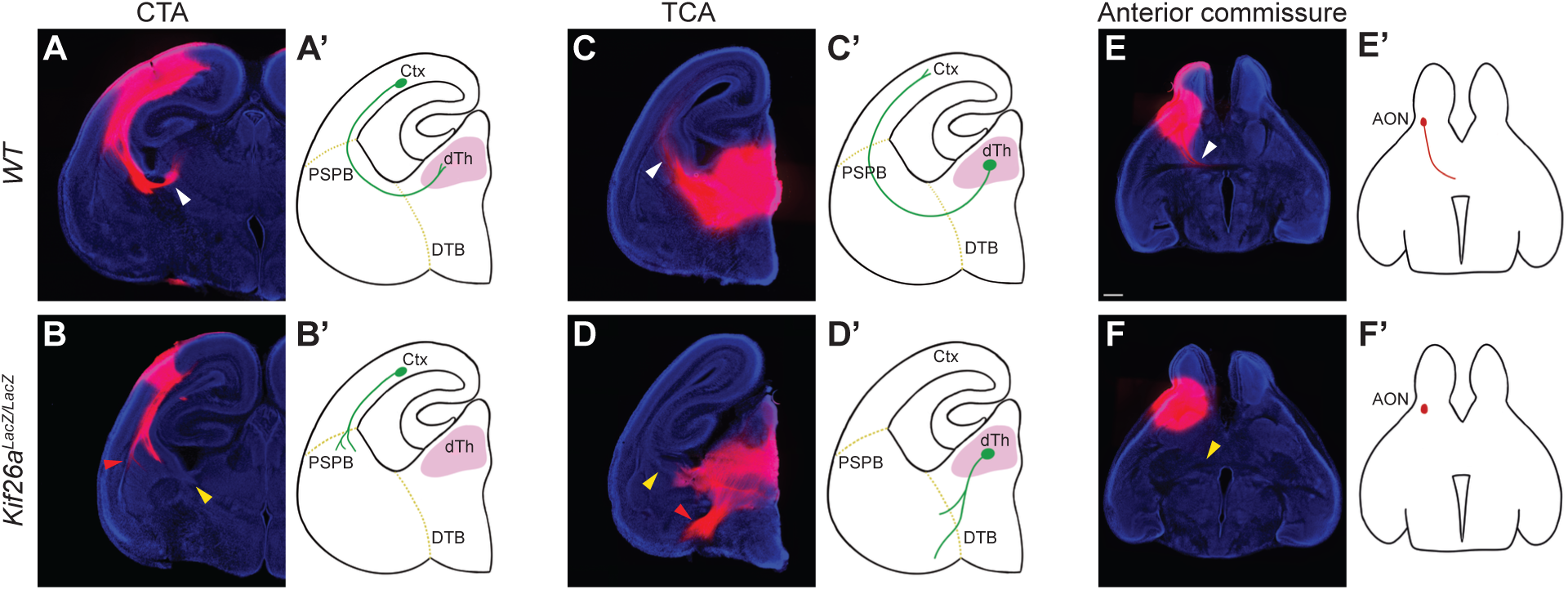
Axon outgrowth and guidance defects in the *Kif26a^LacZ/LacZ^* brain. **(A, B)** DiI crystals were placed in the somatosensory cortex of E18.5 brains to trace the corticothalamic axons (CTA) in WT (**A**) and *Kif26a^LacZ/LacZ^* (**B**) brains. (**A’, B’**) Schematics illustrating the trajectories of the CTA in WT (**A’**) and *Kif26a^LacZ/LacZ^* (**B’**) brains. (**C, D**) DiI crystals were placed in the dorsal thalamus (dTh) of E18.5 brains to trace the thalamocortical (TCA) axons in WT (**C**) and *Kif26a^LacZ/LacZ^* (**D**) brains. (**C’, D’**) Schematics illustrating the trajectories of the TCA in WT (**C’**) and *Kif26a^LacZ/LacZ^* (**D’**) brains. The normal destinations of the analyzed tracts are marked with white and yellow arrowheads in WT and mutant samples, respectively. The red arrowheads mark the misrouted axons. Ctx, cortex; PSPB, pallial-subpallial boundary; DTB, diencephalon-telencephalon boundary. Scale bar: 500 µm

We also traced the anterior branch of the AC (ACa) by placing DiI crystals in the anterior olfactory nucleus (AON). In the WT brain, ACa fibers coursed caudally and then medially to cross the midline, with some labelled fibers entering the olfactory bulb (**Fig. 4E, 4E’**). In *Kif26a^lacZ/LacZ^* brains, only diffused local staining was detected (**Fig. 4F, 4F’**). Thus, loss of *Kif26a* either prevents ACa outgrowth or causes an early guidance error that is followed by axon retraction or death.

### Kif26a functions cell autonomously to control axon development and guidance

Axon pathfinding depends on interactions between the growing axons and guidance cues presented along its trajectory. To test whether Kif26a acts cell autonomously within the projection neurons to guide their axons, we used the *Emx1^Cre^* driver to selectively attenuate *Kif26a* expression in neurons of the dorsal telencephalon (Gorski et al., 2002). L1-CAM staining of *Kif26a^f/f^; Emx1^Cre^* brains revealed a substantial loss of the anterior branch of the anterior commissure, as well as a reduction in the cross-sectional area of internal capsule fibers, consistent with a cell-autonomous role for Kif26a in axon guidance (**Fig. 5B, 5B’, C, E**). Unlike the germline *Kif26a^lacZ/LacZ^* mutants, however, the number of internal capsule fibers was unaffected in *Kif26a^f/f^; Emx1^Cre^* brains (**Fig. 5D**), an expected result given that *Emx1^Cre^* is not active in the thalamus and therefore spares TCA contribution to the internal capsule.

**Figure 5.**
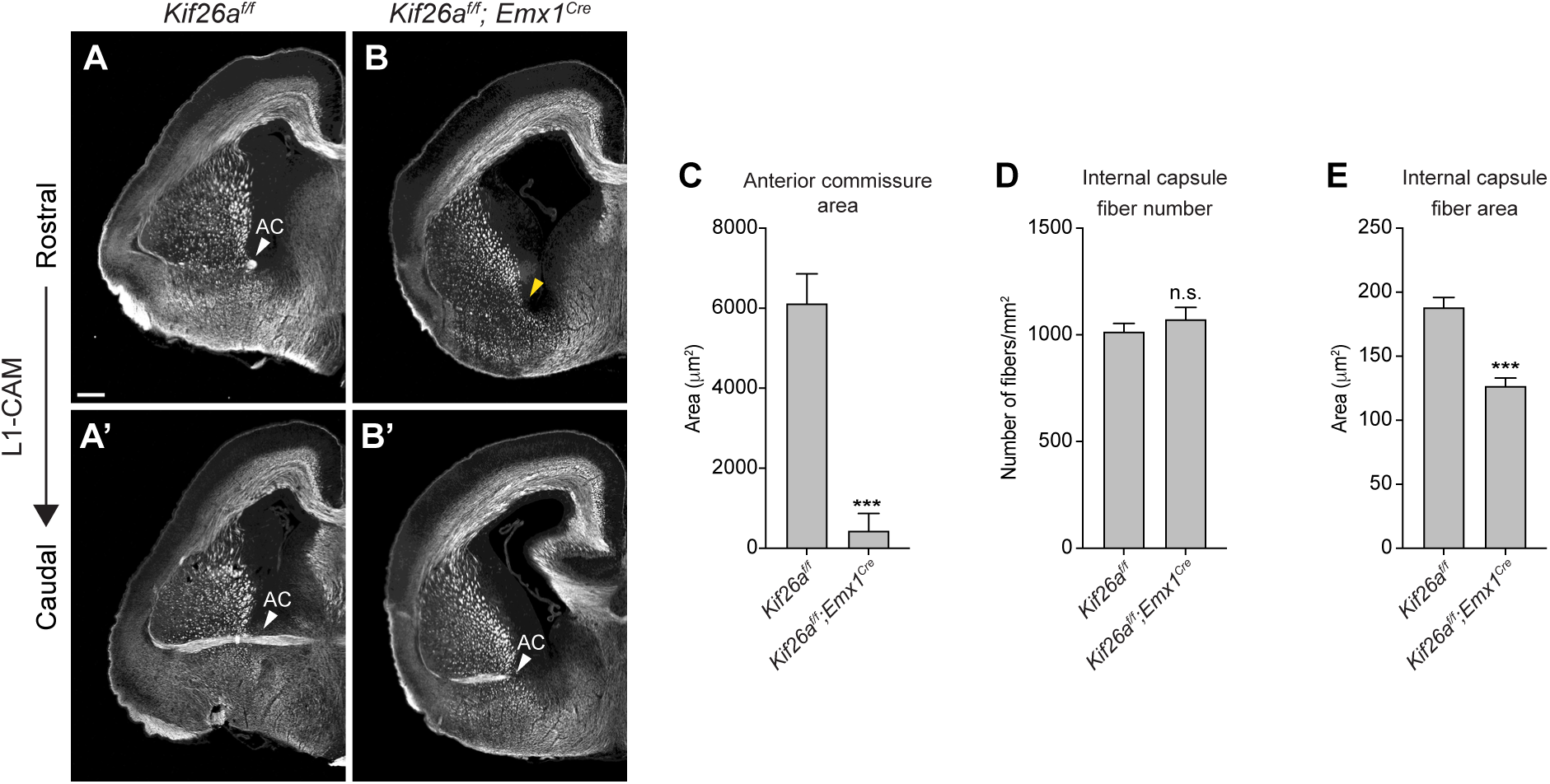
Cell autonomous role of *Kif26a* in axon outgrowth and guidance. (**A-B’**) L1-CAM immunostaining of E18.5 coronal series showing the normal and compromised axon tracts in *Kif26a^f/f^* (**A, A’**) and *Kif26a^f/f^; Emx1^Cre^* (**B, B’**) brains, respectively. The white arrowheads point to the anterior commissure (AC); the yellow arrowheads point to the absent AC in *Kif26a^f/f^; Emx1-Cre*. (**C-E**) Quantification of the AC and internal capsule phenotypes shown in **B-B’**. Data are represented as mean +SEM of the quantified phenotypes from 3 independent embryos for each genotype. t-Test (unpaired) was performed to determine statistical significance of mutant vs. WT. ***, *p* < 0.001; n.s., not significant. Scale bar: 250 µm.

## Discussion

In this study, we establish Kif26a as a crucial regulator of axon guidance during mouse brain development. Genetic ablation of *Kif26a* causes the misrouting or loss of major axon tracts, including the anterior commissure, thalamocortical and corticothalamic projections of the internal capsule, medial lemniscus, and corticospinal tract. These defects occur without detectable changes in proliferation, apoptosis, or cortical layer organization, indicating that Kif26a’s role in axon guidance is direct. Conditional deletion in dorsal telencephalon recapitulates a subset of these tract defects, confirming that the requirement is cell autonomous.

These findings identify Kif26a as one of the few intracellular proteins with a demonstrated, specific role in axon guidance *in vivo*. This is significant because most well-characterized axon guidance molecules are secreted ligands, cell-surface receptors, or extracellular matrix components, and it remains largely unclear how guidance signals received at the cell surface are transduced within the cell to drive cytoskeletal changes and direct axon behavior. Identifying the intracellular proteins that relay guidance signals has been challenging because these molecules, especially those involved in signal transduction and cytoskeletal regulation, tend to be pleiotropic, and their genetic disruption often produces broad developmental defects that obscure specific guidance phenotypes. Members of the Kif26 family have been linked to cytoskeletal regulation and cell adhesion(Uchiyama et al., 2010, Recuenco et al., 2015, Guillabert-Gourgues et al., 2016, Susman et al., 2017). Our identification of Kif26a as a *bona fide* axon guidance regulator therefore offers a valuable entry point for dissecting how extracellular guidance cues are converted into dynamic changes in axon behavior. An important goal for future work will be to define the cell biological functions of Kif26a during axon pathfinding and to understand how these functions are regulated by guidance signals.

Recent identification of *KIF26A* and *KIF26B* mutations in patients with congenital brain abnormalities underscores the clinical relevance of the Kif26 family (Wojcik et al., 2018, Nibbeling et al., 2017, Qian et al., 2022, Almannai et al., 2023, Akula et al., 2023, Gregg et al., 2024). Prior RNAi experiments in mouse *in utero* electroporation and human organoid models showed that *Kif26a* knockdown can cause apoptosis, and impair radial migration and axon/dendrite outgrowth (Qian et al., 2022); our findings reveal axon misguidance as a previously unrecognized pathogenic mechanism underlying the patient phenotypes. It will be important to determine whether patients carrying *KIF26A* mutations display tract defects similar to those observed in *Kif26a* mutant mice in future studies. Given that *KIF26B* mutations are independently associated with cerebellar and spinocerebellar pathology, defining the neurodevelopmental functions of Kif26b warrants further investigation.

An important phenotypic difference between humans and mice is that agenesis of the corpus callosum, which is highly prevalent among patients with *KIF26A* mutations but absent in *Kif26a* mutant mice. One possible explanation is that corpus callosum formation in humans is more sensitive to genetic perturbations (Edwards et al., 2014, Suarez et al., 2014). Human callosal axons are more numerous, traverse a greater distance, and encounter more complex tissue environments than in mice. Thus, considerably more genes are linked to corpus callosum defects in humans than in mice (Kamnasaran, 2005). Another intriguing possibility is that the functions of KIF26A may have expanded during evolution of the human brain (Suarez et al., 2014). Our expression analysis showed that in mice, *Kif26a* is largely restricted to the deeper layers (V and VI) from which the evolutionarily ancient tracts, such as the anterior commissure and internal capsule, originate. Correspondingly, these are the tracts most strongly affected in the *Kif26a* mutant brain. In contrast, *KIF26A* expression in humans appears to have expanded to the upper cortical layers, particularly layer IV, which contributes the majority of callosal fibers, consistent with the specific requirement for *KIF26A* in corpus callosum formation in humans (Fame et al., 2011, Qian et al., 2022). It is unclear why RNAi-based models (Qian et al., 2022), but not the *Kif26a* knockout mice, produced the radial migration and corpus callosum phenotypes, though differences between acute knockdown and constitutive ablation may be a contributing factor.

The signaling pathway through which Kif26a controls axon guidance remains to be determined. However, many of the fiber tract defects observed in *Kif26a* mutant mice phenocopy those seen in mice carrying mutations in *Fzd3*, *Celsr3*, or *Dystroglycan* — three components of a conserved signaling pathway that regulates axon guidance (Wang et al., 2002, Tissir et al., 2005, Lindenmaier et al., 2019). Combined with the well-documented genetic interactions between Vab-8 and Wnt/Fzd signaling in *C. elegans* (Zinovyeva et al., 2008, Pan et al., 2006, Jain and Lundquist, 2025), these observations raise the possibility that Kif26a participates in the Fzd3-Celsr3-Dystroglycan axis, a hypothesis that should be experimentally tested in the future.

Taken together, our findings and the rapidly growing body of literature demonstrate the importance of the Kif26 family in nervous system development, a role conserved from *C. elegans* Vab-8 to mammalian Kif26 proteins. A deeper understanding of this family will not only reveal new principles governing neural circuit formation but also shed light on the mechanisms underlying neurodevelopmental disorders.

## Materials and Methods

### Antibodies

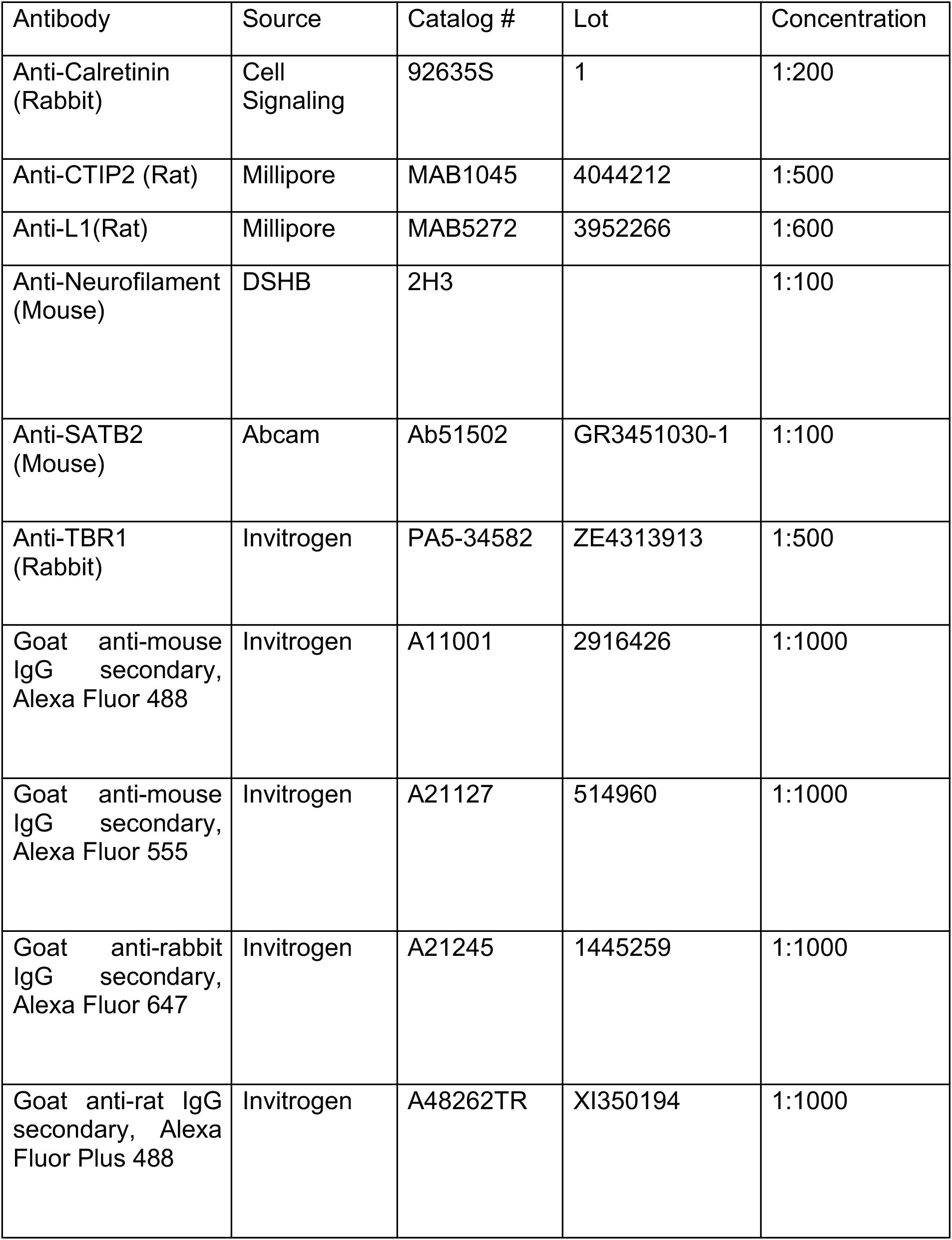
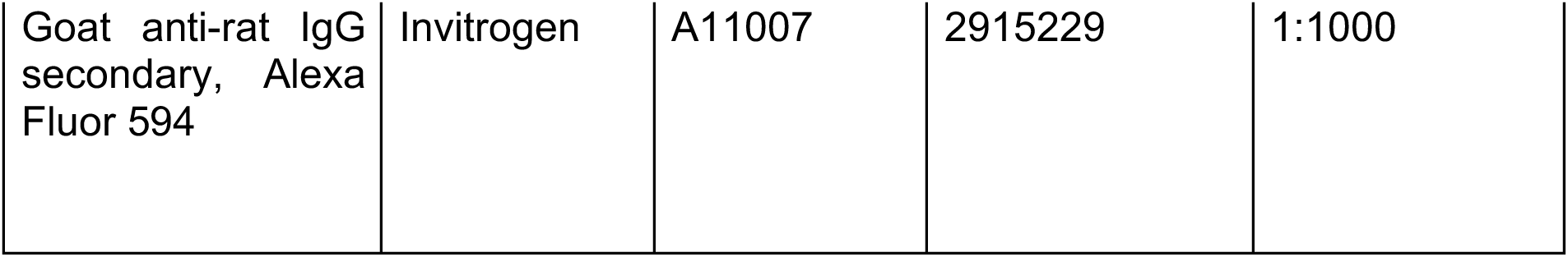

| Antibody | Source | Catalog # | Lot | Concentration |
| --- | --- | --- | --- | --- |
| Anti-Calretinin (Rabbit) | Cell Signaling | 92635S | 1 | 1:200 |
| Anti-CTIP2 (Rat) | Millipore | MAB1045 | 4044212 | 1:500 |
| Anti-L1(Rat) | Millipore | MAB5272 | 3952266 | 1:600 |
| Anti-Neurofilament (Mouse) | DSHB | 2H3 |  | 1:100 |
| Anti-SATB2 (Mouse) | Abcam | Ab51502 | GR3451030-1 | 1:100 |
| Anti-TBR1 (Rabbit) | Invitrogen | PA5-34582 | ZE4313913 | 1:500 |
| Goat anti-mouse IgG secondary, Alexa Fluor 488 | Invitrogen | A11001 | 2916426 | 1:1000 |
| Goat anti-mouse IgG secondary, Alexa Fluor 555 | Invitrogen | A21127 | 514960 | 1:1000 |
| Goat anti-rabbit IgG secondary, Alexa Fluor 647 | Invitrogen | A21245 | 1445259 | 1:1000 |
| Goat anti-rat IgG secondary, Alexa Fluor Plus 488 | Invitrogen | A48262TR | XI350194 | 1:1000 |
| Goat anti-rat IgG secondary, Alexa Fluor 594 | Invitrogen | A11007 | 2915229 | 1:1000 |

### Mouse strains

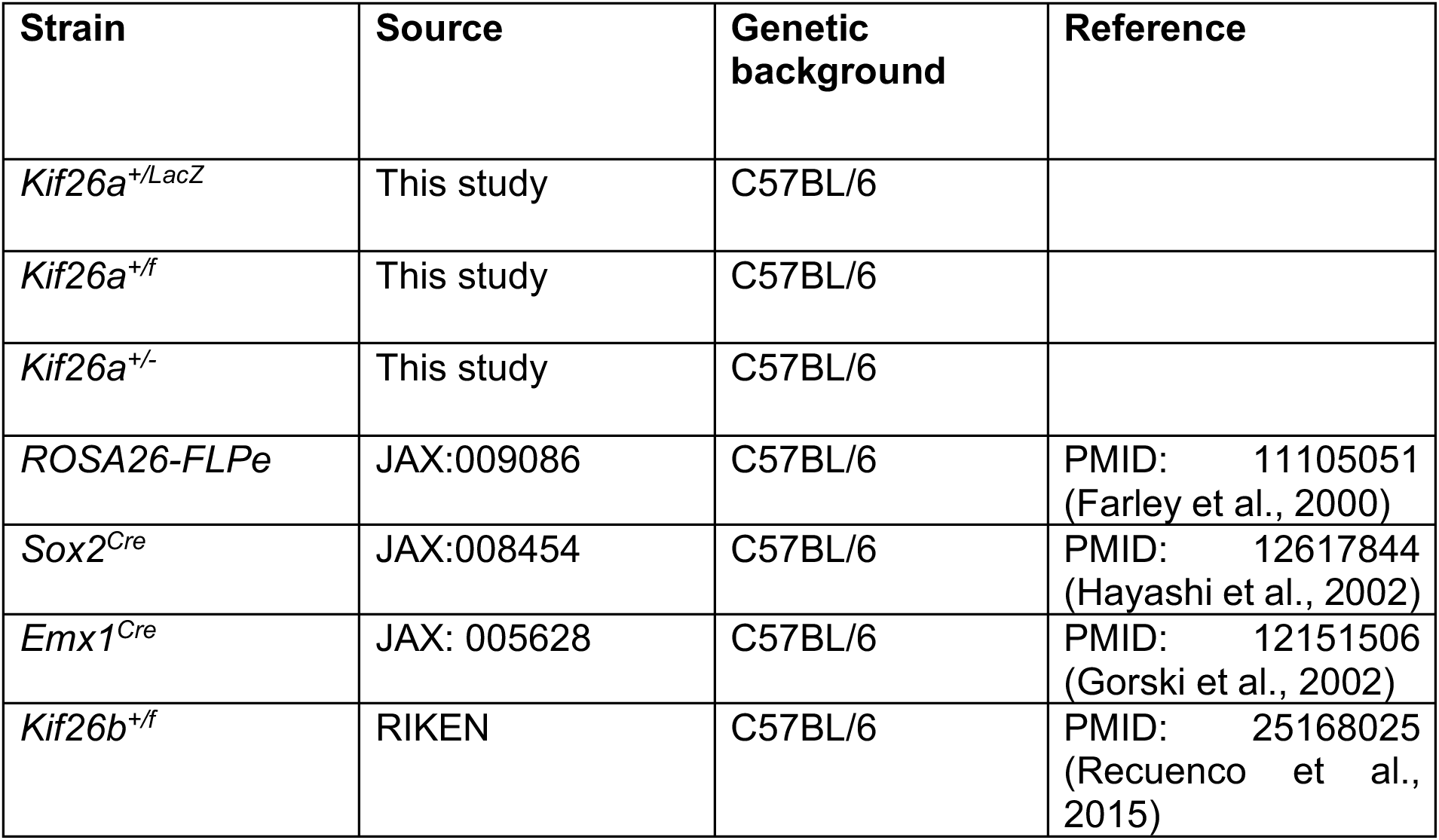

### Animal work

Mice used in the study are listed in the table above. *Kif26b+/-* mice were produced by crossing *Kif26b^f/f^* mice ((Recuenco et al., 2015), stock # CDB0800K, RIKEN CDB, Japan, http://www2.clst.riken.jp/arg/mutant_list_file/CDB0800K.html) to *Sox2^Cre^*. All experiments involving the use of animals were conducted according to protocols and guidelines approved by the institutional animal care and use committee at the University of California, Davis. For timed mating, the morning of plug appearance is defined as embryonic day 0.5 (E0.5).

### In situ hybridization/RNAscope

E14.5 embryo heads were fixed with 4% PFA in PBS at 4°C for 16-18 hours. Samples were washed in PBS, equilibrated in 30% sucrose in PBS for at least 18 hr, and then embedded in a 1:1 mixture of OCT and 30% sucrose in PBS. 20μm thick coronal sections were cut using a cryostat and collected on SuperFrost Plus glass slides. Probes (C1 channel) for *Kif26a* and *Kif26b* were custom designed and purchased from Bio-Techne. In situ hybridization and signal detection were performed according to the manufacturer’s instructions using the RNAscope 2.5 HD Reagent kit - BROWN (Bio-Techne, #322300). Briefly, sample slides were heated in a hybridization oven at 60°C for 30 min followed by incubation in cold 4% PFA in PBS for 1hr. Sample slides were then dehydrated in a 50%, 70% and 100% ethanol series for 5 min in each ethanol concentration at room temperature (RT). After drying the samples for 5 min at RT, they were treated with hydrogen peroxide provided with the kit for 10 min at RT. Sample slides were then incubated in boiling antigen retrieval solution provided with the kit for exactly 5 min, washed 2x in water, dehydrated in 100% ethanol by moving the slides up and down 3-5 times and air dried for 5 min. A barrier around the cryosections was created using a hydrophobic barrier pen. The barrier was allowed to dry for 20 min. The sections were incubated with Protease Plus at 40°C for 30 min. The sample slides were washed 2x in wash buffer (1x concentration, diluted from the 50x stock provided with the kit) and stored in 5x SSC buffer overnight.

The next day, sections were incubated with AMP1 to 6 solutions with alternating incubations of 30 min and 15 min. All incubations were done at 40 °C in the HybEZ oven. Sample slides were washed 2x in fresh wash buffer (1x concentration) between each AMP incubation. To develop the signal, equal volumes of BROWN A and BROWN B solutions were mixed to make the DAB solution. Sections were incubated in the DAB solution at RT for 10 min and washed 2x with distilled water. Sections were then counterstained with hematoxylin, dehydrated in 70% and 95% ethanol for 5 min each, incubated in xylene for 5 min and mounted using Permount (Fisher Scientific, #SP15-100). Sections were then imaged on an Evident APX100 microscope using a UPLXAPO 20X objective.

### Analysis of scRNA-seq data

Single-cell RNA-sequencing data analyses were performed using previously published datasets from E14.5 and E18.5 WT mouse embryos(Loo et al., 2019, Wittmann et al., 2021). All analyses were conducted using RStudio (v4.5.1) and the Seurat R package (v5.3.1), with a consistent random seed maintained across all analyses for reproducibility. The E14.5 dataset consisted of a single count matrix, while the E18.5 dataset was composed of four individual replicates that required subsequent integration.

Individual Seurat objects were created from the raw count matrices. Cell-specific metrics, including the percentage of mitochondrial transcripts (*percent.mt*) and ribosomal transcripts (*percent.ribo*) were calculated. QC filtering thresholds were adopted directly from the original publications to ensure consistency:

E14.5: UMI>500 and <10% of mitochondrial transcripts

E18.5: 1000-5000 genes per cell, 1800-10,000 UMI and <6% of mitochondrial transcripts

To correct for potential batch effects among the four E18.5 replicates, the data were integrated using the SCTransform-based integration workflow in Seurat. Integration features *(nfeatures = 3000*) were selected (*SelectIntegrationFeatures*), integration anchors were identified (*FindIntegrationAnchors*), and the datasets were merged using *IntegrateData*, creating a batch-corrected “integrated” assay for downstream analysis. The E14.5 dataset was processed directly using the *SCT* assay. Dimensionality reduction and clustering were performed on the normalized assay for each sample (SCT for E14.5, and integrated for E18.5). Principal Component Analysis (*RunPCA*) was performed. 30 PCs were used for both datasets to generate the UMAP visualization (*RunUMAP*). A shared-nearest neighbor graph was constructed (*FindNeighbors*), and the Louvain algorithm (*FindClusters*) was used to identify clusters with specific resolutions: 0.9 for E14.5 and 0.8 for E18.5. To visually separate an identified population in the UMAPs, the coordinates for Cluster 20 were manually shifted by x = -4.5 on the final plot of the E14.5 data and Cluster 20 and 18 were shifted by x = -4.5 and y = 3, respectively, in the E18.5 data, creating a “*umap_shifted*” reduction in both analyses. Cluster identity was defined using known marker expression signatures as previously described using the *FeaturePlot* and *DotPlot* functions (Shekhar et al., 2016). Feature Plots were generated for *Kif26a* and *Kif26b* on the UMAP reduction. Analysis was performed by categorizing cells based on whether their normalized SCT expression value was greater than 0.

### Genetic modification of the mouse *Kif26a* locus

The *Kif26a* locus was genetically modified via homologous recombination in mouse embryonic stem cells. The *Kif26a* targeting construct was designed based on the general strategy described by Skarnes *et al*. (Skarnes et al., 2011) and built in-house using a PGK-neo-DTAII plasmid (generated in (Soskis et al., 2012)), as the backbone. The construct contained *lox2272* sites, a mutant *loxP* variant, to avoid potential cross-recombination with standard *loxP* sites. The construction entailed the following key steps: 1) a *lox2272* sequence was inserted between the NheI and HindIII sites; 2) a fragment containing exon 4 of *Kif26a* was amplified from the genomic DNA of C57BL/6 mice by PCR, cloned, sequence verified and inserted into the NheI site. An FseI site was introduced near the 5’ end of this exon 4-containing fragment; 3) a fragment containing the 4.7 kb 3’ homology arm was PCR amplified from C57BL/6 DNA, cloned, sequence verified and inserted into the HindIII site via Gibson assembly; 4) a fragment containing the 4.4 kb 5’ homology arm was similarly PCR amplified from C57BL/6 DNA, cloned, sequence verified, and inserted between the PacI and SacII sites by Gibson assembly. An AflII site was added at the 5’ end of this fragment for subsequent linearization of the final targeting construct prior to ES cell transfection; 5) a fragment containing the Neo cassette followed by *frt* and *lox2272* was PCR amplified from the L1L2_Bact_P vector (purchased from KOMP), cloned, sequence verified, and inserted into the targeting construct between the AscI and NheI sites via restriction digest and ligation; 6) a final fragment containing *frt*, splice acceptor, *IRES-LacZ*, polyA signal, and *lox2272* was PCR amplified from the L1L2_Bact_P vector, cloned, sequence verified, and inserted into the targeting construct between the PacI and AscI sites via restriction digest and ligation. Successful assembly at each step was confirmed by restriction digest and Sanger sequencing.

The final targeting construct, 21.3 kb in size, suppresses *E. coli* growth in liquid culture. Therefore, *E. coli* harboring the construct were grown as a lawn on Luria-Bertani agar plates and collected by scraping for plasmid DNA purification. The targeting construct was linearized by AflII restriction digest, purified by ethanol precipitation, and electroporated into albino C57BL/6 ES cells. G418-resistant colonies were picked, expanded, and screened initially by PCR to detect successful recombination of both 5’ and 3’ homology arms into the Kif26a locus. Clones that were positive in the PCR screens were further confirmed by Southern blotting. The targeting efficiency was 39%. Two validated ES cell clones were injected into blastocysts. Chimeric mice derived from the injected blastocysts were bred, and offspring were assessed for germline transmission of the targeted allele.

### X-gal staining

For E14.5 embryos, brains were dissected out and fixed with 2% PFA, 0.2% glutaraldehyde in PBS for 30 min at 4°C followed by 3 washes in PBS at RT. For cryosectioning, brains were equilibrated in 30% sucrose in PBS overnight and embedded in a 1:1 mixture of OCT and 30% sucrose in PBS. 20μm thick coronal sections were cut using a cryostat and collected on SuperFrost Plus glass slides. To perform X-gal staining, fixed whole brains or sections were permeabilized in 0.02% Igepal, 0.01% sodium deoxycholate, 2mM magnesium chloride, and 0.1M sodium phosphate buffer pH 7.5 3x for 15 min each, followed by immersion of the samples in X-gal staining solution (0.5mg/ml X-gal, 5mM potassium ferricyanide, 5mM potassium ferrocyanide, 0.02% Igepal, 0.01% sodium deoxycholate, 2mM magnesium chloride, and 0.1M sodium phosphate pH 7.5) overnight. Samples were then washed 3x in PBS for 5 min each, post-fixed in 4% PFA in PBS for 30 minutes at RT, and washed 2x in Milli Q H_2_O. Sections were counterstained in nuclear fast red solution (G Biosciences, #786-1054) for 5 sec, mounted with Fluoromount-G without DAPI, and imaged on an Evident APX100 microscope using UPLXAPO 10X objective. Whole brains were postfixed and stored in 4% PFA until imaging.

### Reverse transcription and qPCR

Total RNA was isolated from E10.5 or E12.5 embryos using the TRIzol reagent (Thermo Fisher, #15596026) following manufacturer’s instructions. cDNA was synthesized from 2.5μg of total RNA per 10 μl reaction volume using Maxima H minus reverse transcriptase (Thermo Fisher, #EP0751) and oligo (dT)_18_ primers. Two reactions were set up and incubated at either 50 °C or 65 °C for 1 hr. After the reactions were terminated by heating at 85 °C, the cDNA generated at the two different temperatures were pooled and used as the template for qPCR. qPCR was performed using QuantiNova SYBR Green PCR Kit (Qiagen, #208054) on a StepOnePlus Real-Time PCR system (Applied Biosystem). The following primers were used:

*Mouse Kif26a* forward GAGAGCCACAGGTCTACAGCGG

*Mouse Kif26a* reverse CTGGGAGGGCTTGGCTTCATTC

*Mouse Gapdh* forward AGGTCGGTGTGAACGGATTTG

*Mouse Gapdh* reverse TGTAGACCATGTAGTTGAGGTCA

Relative expression levels were normalized to *GAPDH* using the mean of the comparative Ct (2^−ΔΔCt^) values for each sample group.

### Immunohistochemistry

Embryonic brains or whole heads were dissected and fixed with 4% PFA in PBS at 4°C for 16-18 hours. Samples were washed in PBS 3 times at RT, equilibrated in 30% sucrose in PBS overnight at 4°C and embedded in a 1:1 mixture of OCT and 30% sucrose in PBS. 20μm thick coronal sections were cut using a cryostat and collected on SuperFrost Plus glass slides. Sections were blocked with 5% normal goat serum and 0.3% Triton X-100 in PBS (PBS-T) for 1 hr, followed by incubation with diluted primary antibody solution in the blocking solution at 4°C overnight. Slides were washed 5x with PBS-T and then incubated for 1 hr with fluorescently labelled secondary antibodies diluted in blocking solution. Sections were washed 5x in PBS-T and 3x in PBS, mounted using Fluoromount G with DAPI (Southern Biotech, #0100-20) and imaged on an APX100 microscope using UPLXAPO 20X objective.

Quantification of axon tract phenotypes was done using Fiji. The area of the anterior commissure was measured in the region just before it turns towards the midline. The number and area of the fibers in the internal capsule were analyzed using automatic thresholding and the “analyze particles” feature in Fiji. The thickness of the intermediate zone was quantified by measuring the thickness of L1-CAM positive fibers in the somatosensory cortex. The upper and lower boundaries of the L1-CAM positive fibers were marked and a line perpendicular to the boundaries was drawn. The length of this line was defined as the thickness of the intermediate zone in the somatosensory cortex.

The cell distribution in the cortex (**Fig. 3 Supplement 1 panel A**) was analyzed as previously described using RapID software (Sekar et al., 2021). Briefly, a rectangular region of interest (ROI), divided into 8 equally sized bins, was manually placed on the cortex to encompass the cortical plate. The upper boundary of the ROI (bin 1), set using the DAPI channel, was placed immediately superficial to the marginal zone. The lower boundary of the ROI (bin 8), set using the TBR1 channel, was placed at the lower limit of the TBR1-positive layer (the layer VI/subplate border). Subsequently, the length of the ROI (the distance between the DAPI-defined upper and TBR1-defined lower boundaries) was extended by 10% on both the upper and lower boundaries to ensure that cells located at or near the boundaries were included in the analysis. The number of cells positive for SATB2, CTIP2, TBR1 in each of the 8 bins was quantified using the software. For samples with cortices that exhibit significant curvature, multiple adjacent, narrower ROIs were drawn side-by-side to ensure that the upper and lower boundaries could be accurately defined; the dimensions of these adjacent, narrower ROIs were kept consistent to minimize bias. The cell counts for each marker (SATB2, CTIP2, TBR1) within the corresponding bins of these adjacent ROIs were summed to yield a final, composite cell count for that specific bin across the entire curved region. The final graph was plotted as percentage of cells positive for each marker in the individual bins.

The thickness of each cortical layer (**Fig. 3 Supplement 1 panel C**) was determined by marking the upper and lower boundary of the positive cells in each marker and measuring the distance between the two boundaries. The cell density (**Fig. 3 Supplement 1 panel D**) was determined by counting the number of cells positive for the marker of interest in a defined area and represented as cells/mm^2^.

### EdU labeling

Pregnant dams were injected intraperitoneally with 25mg EdU (APEXbio, #B8337) per kg body weight on gestational day 12.5. Embryos were harvested 4 h later, and the whole heads were collected and processed for cryosectioning as described above. Detection of EdU-labelled cells was performed using the Click-iT EdU cell proliferation kit (Invitrogen, #C10338,) following the manufacturer’s instructions. Images were taken on an APX100 microscope using UPLXAPO 20X objective and analyzed using RapID. Specifically, an ROI–was drawn over the entire cortical area and the area of the ROI was calculated. Labelled cells were automatically counted by RapID. Data were graphed as EdU-positive cells/μm^2^.

### TUNEL assay

Heads were collected from E14.5 embryos and processed for cryosectioning as described above. 20μm thick coronal sections were cut using a cryostat and collected on SuperFrost Plus glass slides. Detection of apoptotic cells was carried out using the Click-iT™ Plus TUNEL assay kit following the manufacturer’s instructions. Positive control sections were treated with 1 unit of DNAse I (Promega, #M6101) diluted in DNAse I dilution buffer (20 mM Tris-HCl, pH 8.4, 2 mM MgCl2, 50 mM KCl) for 30 min at RT. TUNEL-reacted sections were counter stained with Hoechst 33342, mounted using Fluormount G without DAPI (Southern Biotech, 0100-01), and imaged on APX100 microscope using UPLXAPO 20X objective. Images were analyzed in Fiji using the “analyze particle” function. Thresholding limits were set to a minimum of 68 arbitrary units to identify TUNEL-positive cells. A size threshold was set to include signals with area >3 μm^2^ to exclude background fluorescence. Data were graphed as the number of TUNEL-positive cells/μm^2^.

### Axon tracing using DiI

E18.5 brains were dissected and fixed with 4% PFA in PBS at 4°C for one week. DiI crystals were inserted into the cortex (somatosensory area), thalamus (dorsal area), and anterior olfactory nucleus of the fixed brains. When labeling the thalamus, the brains were hemisected along the mid-sagittal plane to expose the thalamus. Samples were returned to 4% PFA in PBS and incubated at 37°C for 2 weeks (CTA labeling) or 4 weeks (AC and TCA labeling). Brains were then embedded in 3% low melting point agarose in PBS, cut into 100μm thick sections using a vibratome. Sections were countered stained with 1μg/mL DAPI for 15 min and mounted on Superfrost glass slides using Fluoromount G without DAPI. Images were acquired on an APX100 microscope using UPLXAPO 10X objective and extended focal imaging (EFI) projection.

### Statistics

3 biological replicates for each genotype were used in all experiments. Statistical analysis and plotting were performed using GraphPad Prism 10. Additional details specific to individual experiments are described in the respective figure legends.

## Acknowledgements

We thank Jeremy Nathans, James Lauderdale, Sneha Mohan, Tom Glaser, Anna La Torre and Nadean Brown for valuable discussion and guidance. We are grateful for technical support from Sneha Mohan, Brad Shibata, Melia Kaplan, Gen Wen Lim, Miranda Do-Tran, Janelle Santos, Alex Sanchez, Sadhana Vijayanand, Jordan Burrise, Anusha Srinivasan and Keiko Hino. This research was supported by the National Institutes of Health grants 1R35GM119574 and 1R35GM144341 to HHH.

